# An optimized protocol for dual RNA-Seq of human macrophages infected with *Mycobacterium avium*

**DOI:** 10.1101/2021.05.20.443437

**Authors:** Jodie A Schildkraut, Valerie ACM Koeken, Jordy PM Coolen, Reinout van Crevel, Jakko van Ingen

## Abstract

Recently, dual RNA-sequencing (RNA-Seq) has been identified as a powerful tool to study host-pathogen interactions, which is particularly interesting for intracellular pathogens such as mycobacteria. However, due to the complexity of obtaining human host cells, many models rely on the usage of host cells derived from animals or cell lines, which does not accurately mimic the situation in the patient. Furthermore, due to the severe disbalance in host and pathogen RNA content, it is difficult to obtain sufficient sequencing depth for the infecting pathogen. Here, we present an optimized method to perform dual RNA-sequencing on human monocyte-derived macrophages (hMDMs) infected with *Mycobacterium avium (M. avium)*. It is likely that, with slight modifications in multiplicity of infection (MOI) to account for differences in virulence, this methodology will be applicable for other difficult-to-lyse intracellular mycobacteria.

## Introduction

Recently, multiple studies have highlighted dual RNA-sequencing (RNA-Seq) as a powerful tool for studying the dynamics of infection, in particular of intracellular pathogens such as mycobacteria^[1–3]^. While traditional RNA-Seq captures either the transcriptome of the host or pathogen, dual RNA-Seq simultaneously captures both. However, due to the complexity of obtaining human host cells many models rely on the usage of animal host cells or cultured cell lines^[2, 3]^. Furthermore, sufficient sequencing depth of the infecting pathogen is difficult to obtain, due to the severe disbalance between host and pathogen RNA, the latter often only accounting for less than 1% of total RNA. In addition, of the total eukaryotic and prokaryotic RNA, the majority is comprised of rRNA, making mRNA enrichment essential before sequencing can be performed^[4]^. Previous studies have shown that comprehensive characterization of the transcriptional response of bacterial pathogens to the intracellular environment is highly dependent on the number of bacterial reads per samples^[5, 6]^. Even at low numbers of bacterial mRNA reads (40-60 thousand), large changes in transcription can still be identified but smaller responses are missed^[5]^. Ideally, a sequencing depth of at least >10^6^ reads is achieved^[6]^. For samples with relatively low correlation between biological replicates, more reads are necessary and a significant number of differentially expressed genes can be identified with around 2-3 million reads^[5]^.

*Mycobacterium avium (M. avium)* is a nontuberculous *Mycobacterium* (NTM) species and an opportunistic intracellular pathogen in humans^[7]^. Because of *M. avium*’s intricate cell wall, shared by all mycobacteria, mechanical lysis of bacteria for nucleic acid extraction is necessary^[8]^. In this study, we optimized a method for dual RNA-sequencing of human monocyte-derived macrophages (hMDMs) infected with *M. avium.* We believe that, with slight modifications in multiplicity of infection (MOI) to account for virulence of each specific species, this methodology will be applicable for other difficult-to-lyse intracellular mycobacteria.

## Results & discussion

### Optimization of experimental infection protocol

To optimize the MOI used in infection experiments, hMDMs from a single donor were seeded into a 96-wells plate and infected with *M. avium* at a MOI of one, five or ten, as described previously^[9]^. Then, the samples were incubated for 24 hours, after which CFU counting of intracellular bacteria was performed. In addition, to study the effect of removing all extracellular bacteria after one hour on the number of intracellular bacteria after 24 hours, we included samples with and without washing to remove extracellular bacteria at one-hour post-infection. For all included conditions 6 biological replicates were included, with the exception of the MOI 5 condition without washing, in which only three replicates were included due to the limited volume of hMDMs. A MOI 10 condition without washing was also omitted due to the limited volume of hMDMs. The average number of bacteria per macrophage are shown in figure 1A. Our findings show that the number of intracellular bacteria is highest at a MOI of 5 when no washing was performed one-hour post-infection, approximately 10-fold higher than when washing was performed. To ensure no increased cell death resulted from infection with *M.* avium at an MOI of 5, cell viability after 24-hours was determined and no increased cell death was seen, as shown in figure 2B. Therefore, in subsequent experiments a MOI of five, without washing one-hour post-infection, was used.

**Figure 1.**
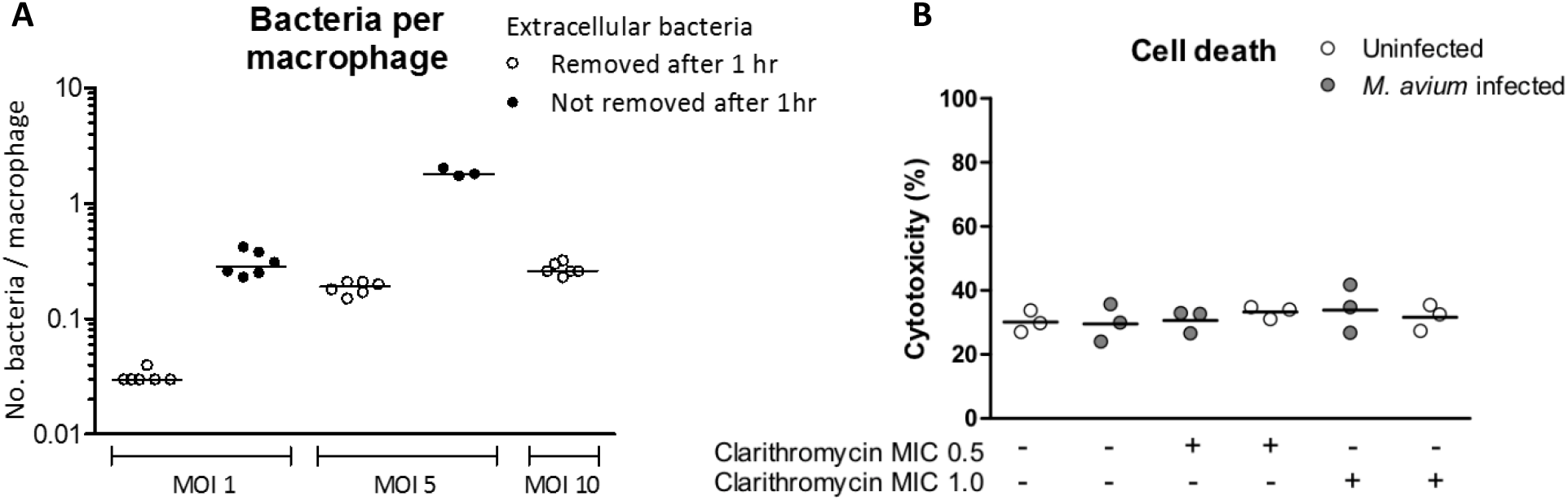
Intracellular *M. avium* count at 24 hours post-infection and survival percentage of macrophages infected with *M. avium* at a MOI of 5 in the presence of differing concentrations clarithromycin. **(A)** The mean number of bacteria per macrophages following 24 hours of infection. For all included conditions 6 biological replicates were included, with the exception of the MOI 5 condition without washing, in which only three replicates were included. **(B)** Macrophage survival after 24 hours of infection at a MOI of 5 in the presence of no, 0.5 or 1 μg/mL clarithromycin.

**Figure 1.**
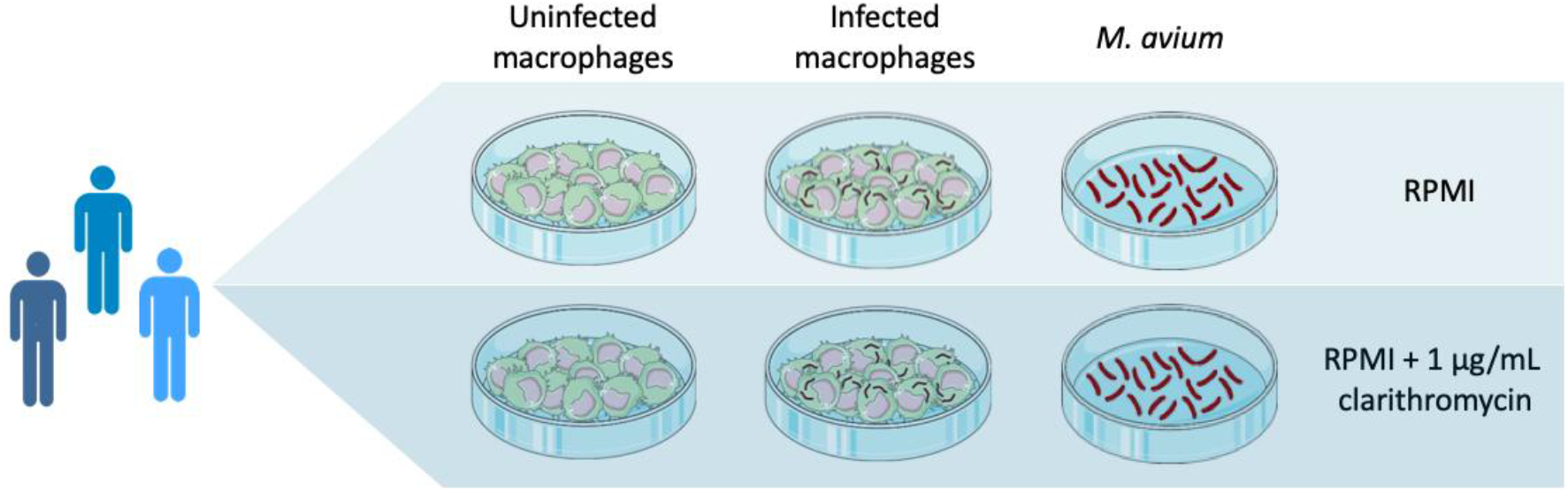
Overview of final experiment set-up for Dual RNA-Seq. Four separate sets of macrophages were cultured from each donor, of which half were infected and half were not, so the comparison could be made between uninfected and infected macrophages. *M. avium* was also included in duplicate so that the comparison could be made between extracellular and intracellular bacteria. In addition, to one of each condition clarithromycin was added at a concentration of 1 μg/mL so the comparison could be made between clarithromycin exposed and non-exposed bacteria.

### Optimization of bacterial RNA enrichment

To circumvent the severe disbalance between *M. avium* and hMDM RNA, we aimed to optimize the enrichment of bacterial RNA. To this end, we exploited the difficult to lyse cell wall of *M. avium*. In a six-wells plate four replicates of 10^7^ hMDMS were seeded and infected with *M.* avium and two replicates of 10^6^ hMDMs were seeded that remained uninfected. After 24 hours of infection, all extracellular *M. avium* were removed by multiple rounds of washing with room temperature (RT) PBS. Subsequently, 3 mL 4M RT guanidinium thiocyanate (GITC) was added to infected macrophages, resulting in lysis of the host cell but not the remaining intracellular *M. avium*. To ensure complete lysis, cells were scraped prior to collection of the complete volume in a 15 mL falcon tube. Samples were then centrifuged at 5000g for 5 minutes at 4°C to pellet all *M. avium*, while ensuring host RNA remained in suspension. Supernatant containing host RNA was then removed by different methods and in different volumes to determine the effect on final RNA concentration as a measure for bacterial mRNA enrichment, results are shown in table 1. Total RNA levels needed to remain high enough for subsequent rRNA depletion (ideally >1000 μg, minimum of >750 ng). Subsequent lysis of *M. avium* was performed by bead-beating prior to RNA isolation and RNA integrity was checked on the TapeStation 2200 (Agilent, Santa Clara, USA) to determine whether enrichment and beating influence RNA quality and ensure all samples had a RIN >8. We then performed library preparation of rRNA-depleted mRNA for infected samples, after which they were pooled equimolar and RNA-Seq was performed to determine the total number of bacterial reads obtained, shown in table 1. Due to the potential risk of dislodging or partially aspirating the *M. avium* pellet we chose decanting of supernatant with a remaining volume of approximately 50 μl for subsequent experiments.

**Table 1.** Overview of bacterial enrichment methods, RNA concentration and total number of bacterial reads

| Sample | Supernatant remaining (µl) | Supernatant removal method | RNA concentration (ng) | Bacterial reads obtained |
| --- | --- | --- | --- | --- |
| Infected 1 | 25 | Pipetting | 685.2 | n/a |
| Infected 2 | 50 | Pipetting | 1295.4 | 1,554,657 |
| Infected 3 | 210 | Decanting | 6670.2 | 434,283 |
| Infected 4 | 80 | Decanting | 2485.8 | 235,386 |
| Uninfected | n/a | n/a | 790.8 | n/a |
| Uninfected | n/a | n/a | 752.4 | n/a |

### Final experimental protocol and enrichment results

To test our finalized protocol, we characterized the transcriptomic response of *M. avium* in the intracellular environment in comparison to that of the extracellular environment. We also studied the effect of the presence and absence of clarithromycin, an antibiotic used in *M. avium* treatment, on all conditions to determine whether the current protocol was also sufficient to differentiate between intracellular conditions. We included hMDMs from three separate healthy donors to account for donor variability. An overview of the experimental protocol is shown in figure 2. An overview of generated reads per condition are shown in table 2. Our findings show that despite differences in the enrichment in the number of bacterial reads, the minimum requirement of >10^6^ reads is achieved in all samples. We then performed principle component analysis (PCA) of all included conditions to determine if a clear transcriptomic profile could be seen within biological replicates of *M. avium* and hMDMs, as shown in figure 3. The PCA plot of all *M. avium* conditions illustrates that in all conditions the biological replicates are highly reproducible and a clear difference in transcriptomic landscape can be seen between each condition. In addition, hMDMs show a reproducible, condition-specific transcriptomic landscape. Interestingly, the PCA plot of hMDMs shows that our current method is able to capture inter-individual variation in the host response to *M. avium* infection.

**Table 2.** Overview of Dual RNA-Seq conditions and generated reads for host and *M. avium.*

| Sample name | Condition | Number of human reads (% total) | Number of *M. avium* reads (% total) |
| --- | --- | --- | --- |
| I1 | Infected macrophages | 68,215,535 (96.6%) | 2,382,532 (3.4%) |
| I2 | Infected macrophages | 33,326,276 (83.3%) | 6,686,911 (16.7%) |
| I3 | Infected macrophages | 27,807,625 (95.8%) | 1,224,237 (4.2%) |
| Ic1 | Infected macrophages + clarithromycin | 53,670,329 (93.3%) | 3,839,453 (6.7%) |
| Ic2 | Infected macrophages + clarithromycin | 49,491,213 (89.1%) | 6,027,656 (10.9%) |
| Ic3 | Infected macrophages + clarithromycin | 29,690,223 (88.0%) | 4,044,943 (12.0%) |

**Figure 3.**
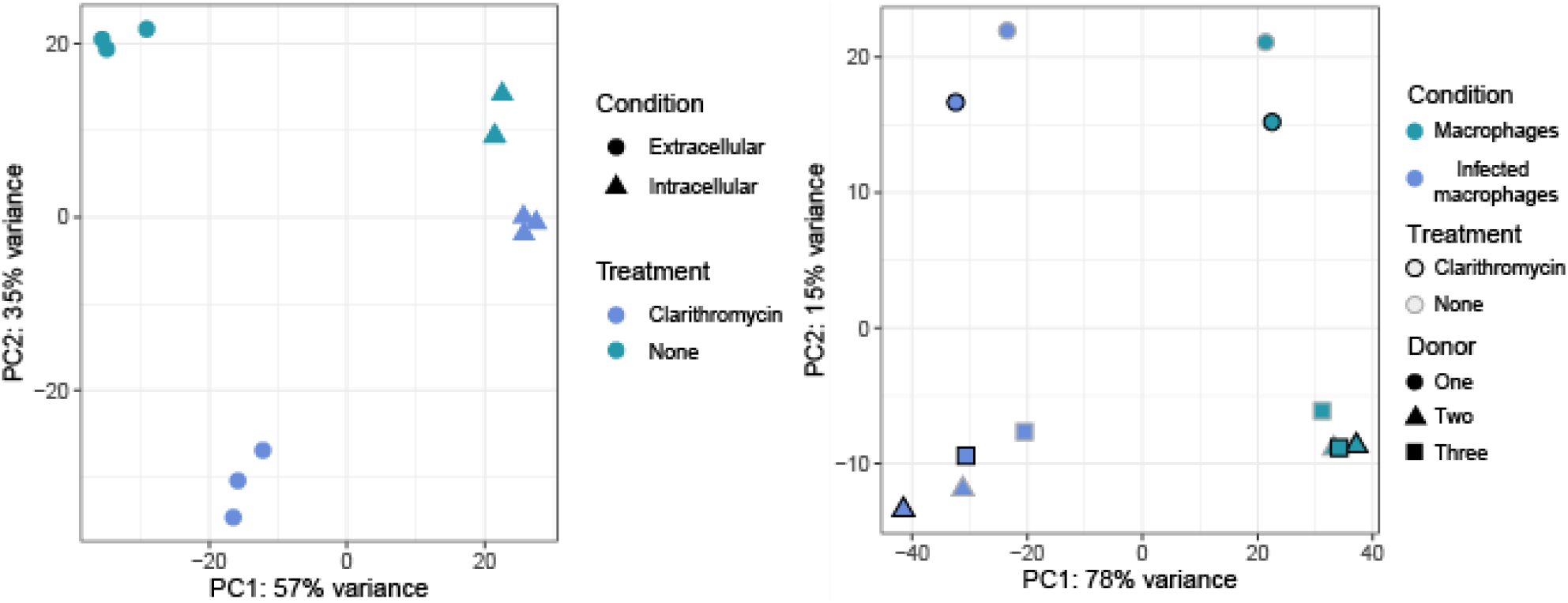
Principle component plot of all conditions. **(A).** PCA plot of all *M.* avium conditions. **(B).** PCA plot of all macrophage conditions.

### Differentially expressed genes in *M. avium*

We then determined whether the achieved sequencing depth was sufficient to study differential gene expression in *M. avium*. To this end we determined the number of differentially expressed genes (DEGs) between intracellular and extracellular *M. avium*, between intracellularly clarithromycin-exposed and intracellular unexposed *M. avium* and between intracellularly clarithromycin-exposed and extracellular *M. avium*. We defined DEGs as genes with a Log2 fold change ≥ 2 or ≤ −2 and a p-value corrected for multiple testing by the Benjamini-Hochberg principle ≤0.05^[10]^. The number of up- and downregulated genes per condition and their overlap is shown in figure 4. A large number of DEGs was seen in intracellular compared to extracellular *M. avium* (516, 304 up- and 212 downregulated). In comparison, in line with the clustering of isolates in the PCA plot, the response to clarithromycin of intracellular *M. avium* was very limited, only 31 DEGs (17 up- and 13 downregulated). Based on the large number of DEGs that were found to be significantly up- or down-regulated we show that the applied dual RNA-Seq protocol was sufficient to study transcriptomic changes in intracellular *M. avium*.

**Figure 4.**
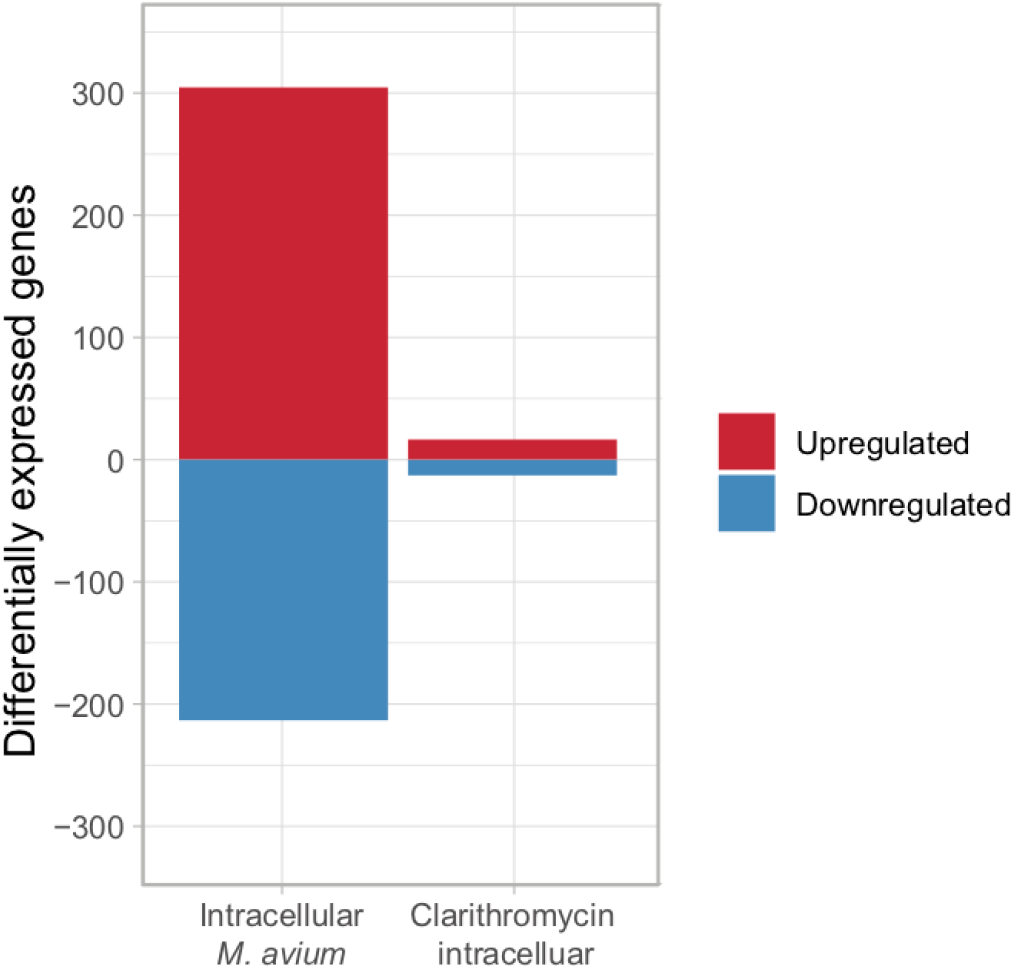
Bar chart depicting number of up- and downregulated genes per condition. When comparing intracellular *M. avium* to extracellular *M. avium* a total of 304 up- and 212 downregulated genes were found. When comparing intracellular *M. avium* clarithromycin-exposed *M. avium* to intracellular unexposed *M. avium* only 17 up- and 13 downregulated genes were found.

### Conclusion

We optimized a dual RNA-Seq method for infection of hMDMs with difficult to lyse mycobacteria, using *M. avium* as a proof of principle. Our findings show that despite the substation disbalance between eukaryotic host RNA and prokaryotic pathogen RNA, an adequate sequencing depth of the intracellular pathogen can be achieved by optimizing MOI, method of infection and mRNA enrichment during RNA isolation. Our findings show that with *M. avium* an optimal ratio of intracellular bacteria to macrophage is achieved with a MOI of 5, and removal of extracellular bacteria only prior to host cell lysis. In addition, we found that pelleting of non-lysed *M. avium* and decanting of host RNA from lysed hMDMs leads to sufficient enrichment of bacterial mRNA. A detailed description of all optimized methods is included in the material & methods section.

## Methods & Materials

### Bacterial strains and antibiotics

The *M. avium* subsp. *hominissuis* ATCC 700898 type strain (synonym: MAC 101; American Type Culture Collection, Manassas, VA) was used for all experiments. Stock vials of the strain were stored at −80°C in trypticase soy broth containing 40% glycerol and were thawed at room temperature prior to each experiment. Clarithromycin was obtained from Sigma-Aldrich and dissolved in methanol.

### Bacterial culture conditions

For macrophage infections, a bacterial inoculum was prepared freshly by allowing them to grow for 5 days in Middlebrook 7H9 broth containing 5% OADC and 0.05% Tween 80 (Sigma-Aldrich, Zwijndrecht, The Netherlands) before making a 0.5 McFarland suspension (+−2,5 * 10^7^ CFU/mL) in PBS. For bacteria-only conditions, bacteria were cultured in 10mL Middlebrook 7H9 broth containing 5% OADC and 0.05% Tween 80 until log-phase and centrifuged at 4000x g for 5 minutes before being resuspended in 10mL RPMI++.

### Differentiation of hMDMs

Healthy volunteers gave their written informed consent for the use of their blood for scientific purposes, as approved by the Ethics Committee of Radboud University Medical Centre, Nijmegen, the Netherlands. Peripheral blood mononuclear cells (PBMCs) were isolated from buffy coats from healthy volunteers (Sanquin Bloodbank, Nijmegen, the Netherlands). Isolation was performed using Ficoll-Paque, involving separation by a density gradient followed by three wash steps in cold PBS and resuspension in RPMI 1640 (Life Technologies, Paisley, UK) supplemented with 10 μg/mL gentamicin, 2 mM GlutaMAX and 1 mM pyruvate. PBMCs were seeded in petri dishes and incubated for 2 hours at 37°C before the non-adherent lymphocytes were washed away with PBS. The adherent monocytes were allowed to differentiate into monocyte-derived macrophages for 6 days at 37°C in culture medium supplemented with 5 ng/mL GM-CSF and 10% human pooled serum. One day prior to the infection, cells were dissociated with Versene and re-seeded in antibiotic-free medium in a 96-well plate (1×10^5^ cells/well).

### Optimization of experimental infection protocol

To optimize multiplicity of infection (MOI), hMDMs were seeded into 24-wells plate at a final concentration of 10^6^/well. *M. avium* bacteria were added at a MOI of 1, 5 and 10. In the MOI 1 and 5 conditions two replicates were included, one in which extracellular bacteria were washed away with PBS after 1 hour and one where they were not. After 24 hours of incubation, cells in all conditions were washed using PBS to remove all extracellular bacteria and cells were lysed using MilliQ. The intracellular bacterial population was then quantified by CFU counting using a 10-fold serial dilution in 0.85% sterile saline. 10 μL per sample of each dilution (including undiluted) were plated in triplicate onto Middlebrook 7H10 agar plates (BD Bioscience) and incubated at 30°C for 3 days before CFU quantification.

### Optimization of Bacterial mRNA enrichment

Following 24 hour of infection, cells infected at a MOI of 5 were washed using RT PBS. Subsequently, 3 mL 4M RT GITC was added to infected macrophages. Cells were then scraped and collected in a 15 mL falcon tube. From all tubes containing infected cells supernatant was decanted leaving differing volumes to enrich for bacterial mRNA, an overview is given in table 1. Samples were transferred to 1.5 mL safe-lock tubes containing one scoop of acid-washed glass beads (<106μM) and 1 mL TRIzol (Invitrogen) containing 20μg/mL GenElute-LPA (Sigma-Aldricht) was added, samples were incubated at RT for 5 minutes and then placed on ice. Samples were then subjected to two rounds of snap freezing in liquid nitrogen followed by bead beating for 30s and 7000 rpm in a MagNAlyser (Roche). Samples were then centrifuged for 1 minute at 13000 g and transferred to new safe-lock tubes. 300 μL chloroform was added to each sample (600 μL to sample MOI 1 due to higher volume) and mixed by hand for 1 minute. Samples were then centrifuged for 5 minutes at 13000g, the aqueous phase was then collected and 600 μL of 70% Ethanol was added to adjust RNA binding conditions. RNA isolation was then performed using the Nucleospin RNA kit (Machery Nagel) according to manufacturers’ protocol from RNA binding onward. RNA integrity was subsequently measured on a TapeStation 2200 (Agilent) and RNA concentration was measured using a NanoDrop (Thermo Scientific). To remove unwanted ribosomal RNA (rRNA) depletion was performed using RiboMinus (Thermo Scientific) human and prokaryote kits. The mRNA library was then constructed using the TruSeq RNA sample preparation V2 kit (Illumina, San Diego, USA) starting from RNA fragmentation. In short, RNA was chemically fragmented prior to cDNA synthesis. End-repair was then performed on constructed cDNA, followed by A-tailing, adaptor ligation and 15 cycles of qPCR. A clean-up of the DNA using AMPure beads was performed between each step. A 1ul aliquot of the library was again run on a TapeStation 2200 to ensure all libraries had the correct length (approximately 290bp). Libraries were pooled equimolar to a final concentration of 4nM and 1.25 pM sample was sequenced in paired-end 2x 75 bp mode on a NextSeq 500 (Illumina) using a High-Output chip.

### Cell viability assay

Macrophage viability during *M. avium* infection at a MOI of 5 in the presence of differing concentrations of clarithromycin was assessed using a LDH cytotoxicity kit according to manufacturer’s instructions (Pierce, Thermo Scientific) as described previously^[11]^. In short, 50 μL of freshly harvested supernatant was mixed with 50 μL LDH reaction mixture in a 96-wells plate and incubated for 30 minutes at RT. The reaction was stopped by the addition of 50 μL stop solution and the plate was measured on a Mithras LB 940 microplate reader (Berthold Technologies) at 485 nm.

### Final experimental protocol

For infected condition hMDMs were seeded into petri dishes containing 10 mL RPMI++ and a final concentration of 10^6^ cells/mL and infected with *M. avium* at a MOI of 5. For uninfected macrophage conditions hMDMs were seeded into a 24-wells plate containing 3 mL RPMI++ per well at a final concentration of 10^6^ cells/mL. For *M. avium* alone conditions, 10 mL of *M. avium* in log-phase growth was collected by centrifugation at 4000x g for 5 minutes and resuspended in 10 mL RMPI++ and seeded into petri dishes. All conditions were completed in sextuplicate and to half of samples from each condition clarithromycin was added at a final concentration of 1 μg/mL (1x MIC), an overview of samples is shown in table 2. Samples were then incubated for 24 hours at 37°C with 5% CO^2^. Following 24h of infection cells were washed using RT PBS, bacterial mRNA enrichment was performed as previously described and RNA isolation was performed. Following RNA isolation, concentration and integrity was checked and mRNA depletion and library preparation was performed using the Illumina TruSeq v2 kit following manufacturer’s instructions from Elute, Prime, Fragment on. Libraries were then pooled to a final concentration of 1.75 nM and sample was sequenced in paired-end 2x 100 bp mode on a NovaSeq 6000 (Illumina, San Diego, sUS) using a S1 chip. A larger chip compared to the previous test experiment was used to generate a larger number of total reads to further deepen the bacterial sequencing and compensate for the larger number of samples in comparison to previous test set.

### Bioinformatic analysis

All obtained reads were mapped to the M. avium 109 (CP029332.1) or the human (Hg38) genome using STAR (v2.7.0)^[12]^. The count table for host samples were generated using STAR (v2.7.0) while *M. avium* count table were generated using the DESeq2 package in R (v3.3.3). Differential expression analysis was performed in R (v3.3.3) using the DESeq2 package^[10]^ and cut-offs were defined as Log2 fold change ≥ 2 or ≤ −2 and a p-value corrected for multiple guessing of ≤ 0.05.

